# Study of Distinctive Diagnostic Characteristics of *Chrysomphalus dictyospermi* and Closely Related Species of *Chrysomphalus* Genus (Insecta: Sternorryncha: Coccoidea)

**DOI:** 10.1101/2020.09.16.295287

**Authors:** N.A. Gura, A.V. Shipulin

**Affiliations:** Researcher of the Research and Methodological Department of Entomology of FGBU “VNIIKR”

**Keywords:** Scales, Chrysomphalus dictyospermi, pygidium, lobes, plates, perivulvar pores, paraphysis

## Abstract

The paper presents the results of microscopic study of the basic diagnostic structures of the pygidium of the Chrysomphalus dictyospermi, which allow to distinguish this species from the closely related ones of the Chrysomphalus genus.

## Introduction

The object of the study was the brown scale insect *Chrysomphalus dictyospermi* (Morgan, 1889)), which belonging to the group of insects (Insecta: Sternorryncha: Coccoidea). By the decision of the Council of the Eurasian Economic Union No. 158 of November 30, 2016, the Unified List of Quarantine Objects of the Eurasian Economic Union had been approved, in which subsequently dictyospermum scale *Chrysomphalus dictyospermi* was included. Earlier this object was included into the lst of quarantine objects that are absent in the Republic of Kazakhstan. This List was put into effect on July 1, 2017 in the territory of all EEU countries. With an increase in the import of planting material for open and protected ground, the risk quarantine pests distribution increases. Detection of quarantine objects on plant products, laboratory research, precise species diagnosis of the object are the main stages of phytosanitary procedures.

**Purpose of this paper —** study of distinctive diagnostic characteristics of dictyospermum scale and closely related species of *Chrysomphalus* genus.

### Aims of the paper

- microscopic study of the female pygidium structure of the *Chrysomphalus dictyospermi* using modern microscopic technologies to identify the main diagnostic features;
- presentation of illustrative material indicating identified diagnostic structures;
- comparative analysis of the diagnostic structures of the *Chrysomphalus dictyospermi* and closely related species: *Chrysomphalus aonidum*, *Chrysomphalus bifasciculatus* and *Chrysomphalus pinnulifer*.

The obtained data will make it possible to reveal distinctive diagnostic features of the dictyospermum scale female from closely related non-quarantine scales species found on similar plant products from distribution countries.

### General information about the pest

*Chrysomphalus dictyospermi* is a polyphage that causes damage to plants of about 80 families (ScaleNet, 2020). Dictyospermum scale feeds on the following forage crops: citrus crops (family: Rutaceae), rose (family: Rosaceae), palms (family: Arecaceae), oil-bearing plants (family: Oleaceae), asparagus (family: Asparagaceae), mulberry (family: Moraceae), legumes (family: Fabaceae), myrtle (family: Myrtaceae), grapes, fruit, subtropical and tropical plants, including potted plants (Cabi, 2020). This scale species is most harmful to citrus and subtropical crops, causing leaf fall, reduced yield and commercial quality of fruits.

## Materials and methods

The diagnostic characteristics of the *Chrysomphalus dictyospermi* female were studied under a microscope (Carl Zeiss Microscopy GmbH). The material for the study was the slides of *Chrysomphalus* sp. from the collection fund of the FGBU “VNIIKR” and scientific collections of plant material with brown shield colonies.

The data of CABI, ScaleNet and scientific publications on *Diaspididae* family were used for the comparative analysis of the diagnostic characteristics of *Chrysomphalus* sp.

## Results

According of the study results, the main diagnostic micro-signs of pygidium of females of the genus Chrysomphalus are the following: the number of lobes, the peculiarities of the structure of plates, the presence of paraphyses and perivulvar pores.

The description of the main diagnostic features characteristic of the shields of the Chrysomphalus genus, as well as the necessary terms and definitions used in the diagnosis of the shields are presented in table 1.

**Table 1.**
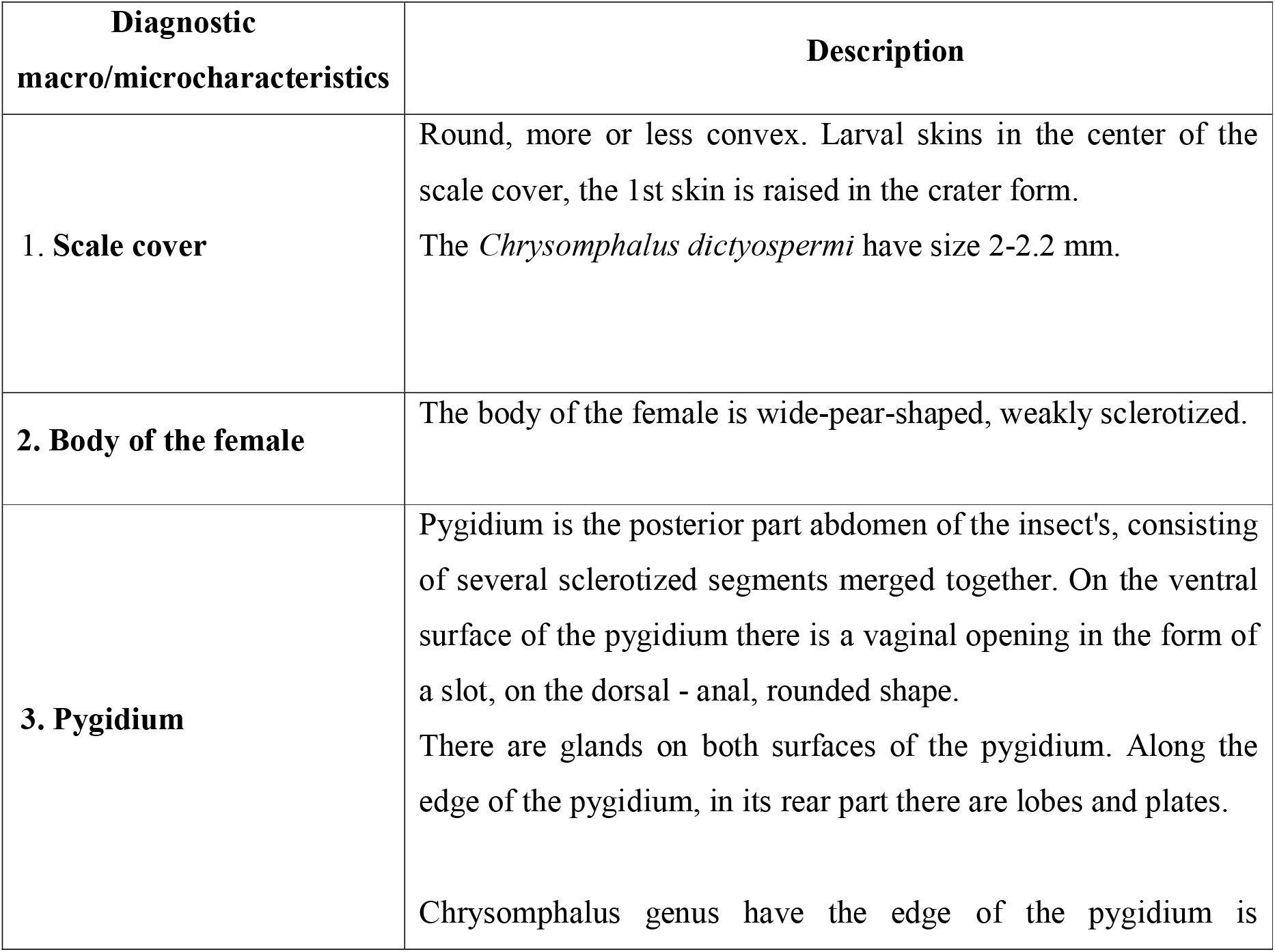
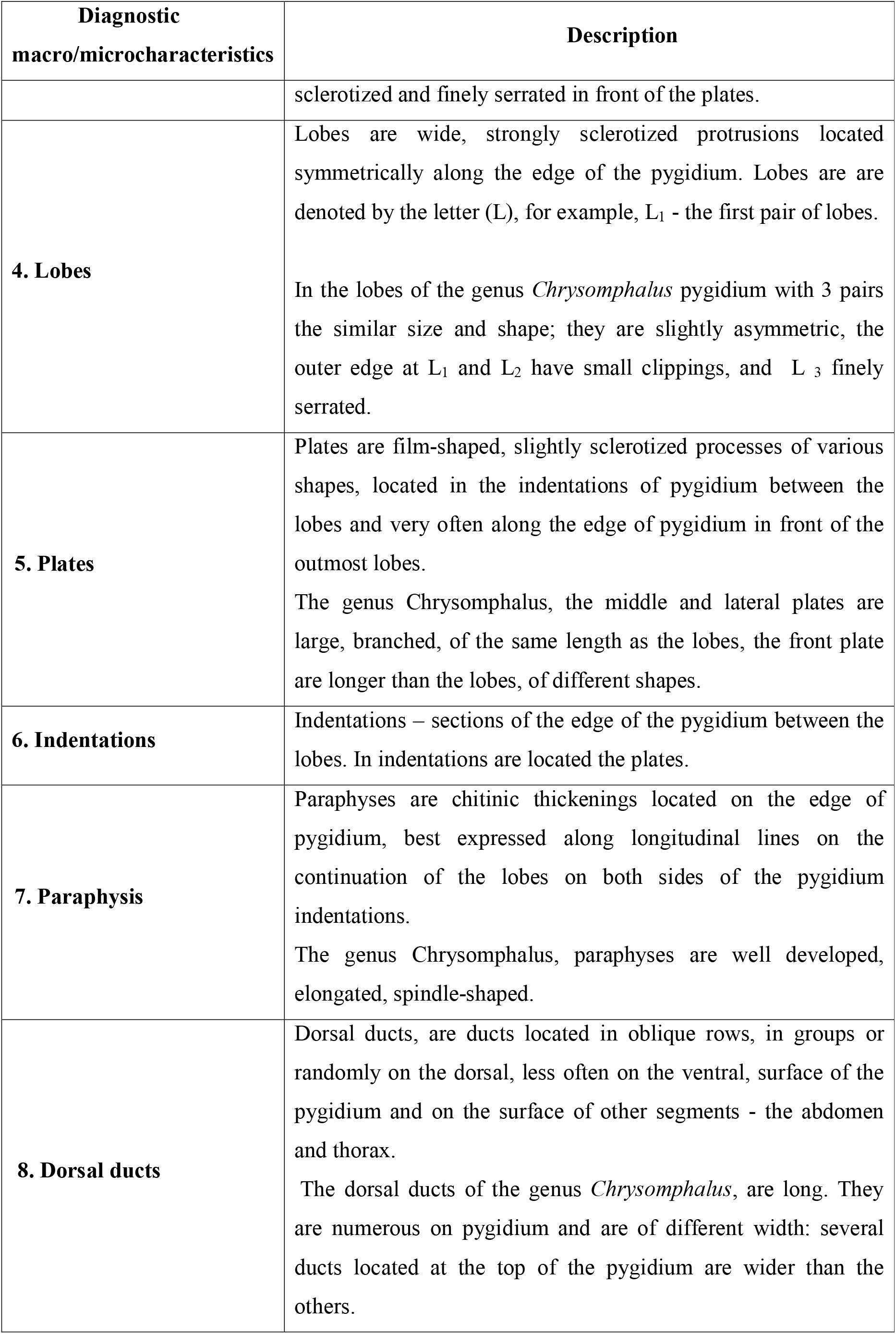
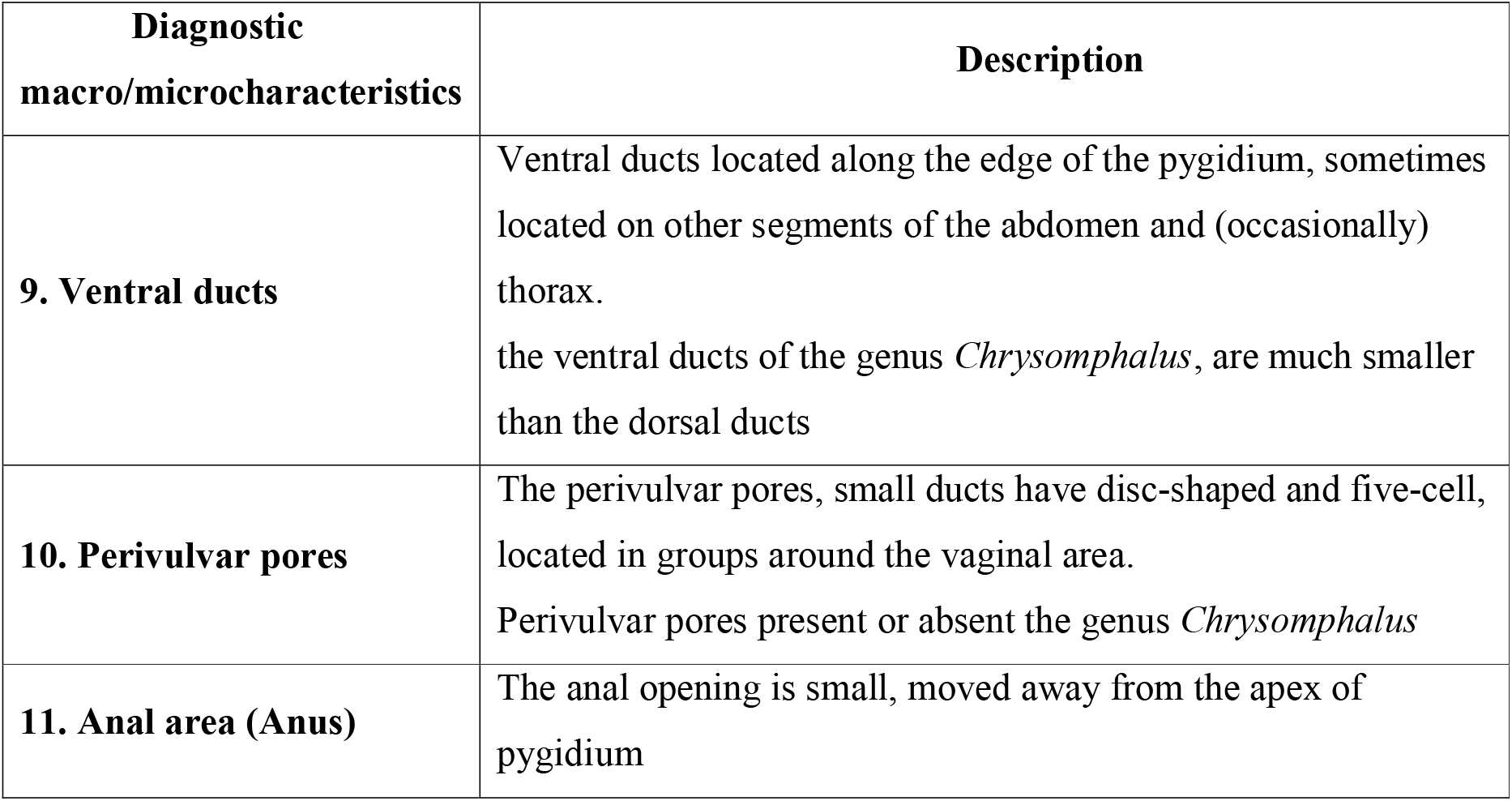
Main terms and description of diagnostic features of Chrysomphalus genus scales (Danzig, 1993)

The main macro signs of the shields of the genus *Chrysomphalus* include the following: the shape and size of the scale cover, its color, number and location of larval skins included in scale cover, the color and shape of the female's body. Macro characteristics of brown shield females are shown in Table 2.

**Table 2.** Description of macro signs of female *Chrysomphalus dictyospermi* (Dantsig, 1993)

| **Diagnostic macrocharacteristics** | **Chrysomphalus dictyospermi** |
| --- | --- |
| Scale cover shape | A weak convex |
| Scale cover size, mm | 1,5-2,0 |
| Scale cover color | Brown with a copper tint |
| Location of larval skins, composing the scale cover | In the center of the scale cover |
| Live female body color | Yellow |
| Body shape | Pear-shaped |

**Fig. 1.**
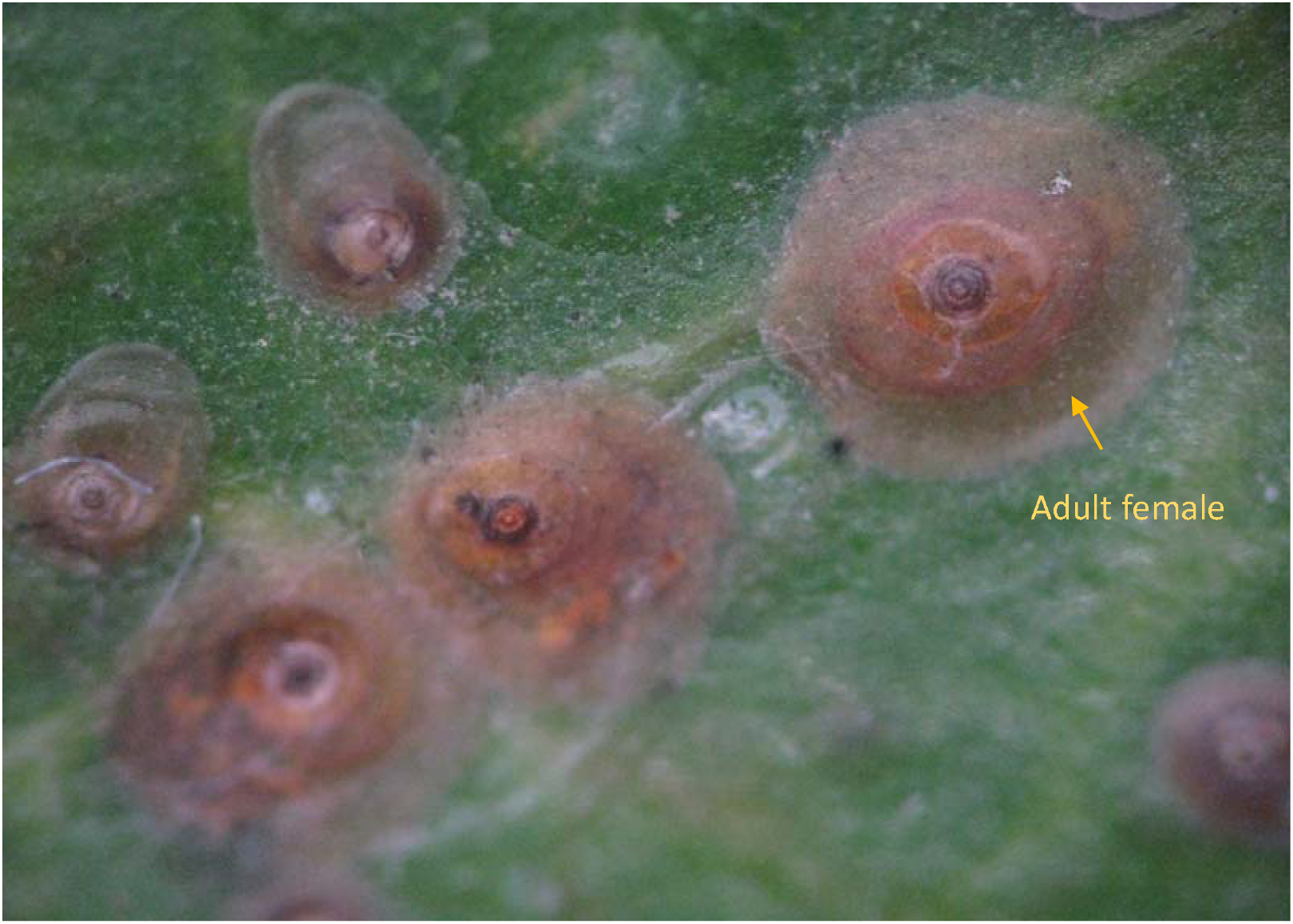
A colony of *Chrysomphalus dictyospermi* (photo authors A.V. Shipulin, N.A. Gura)

**Fig. 2.**
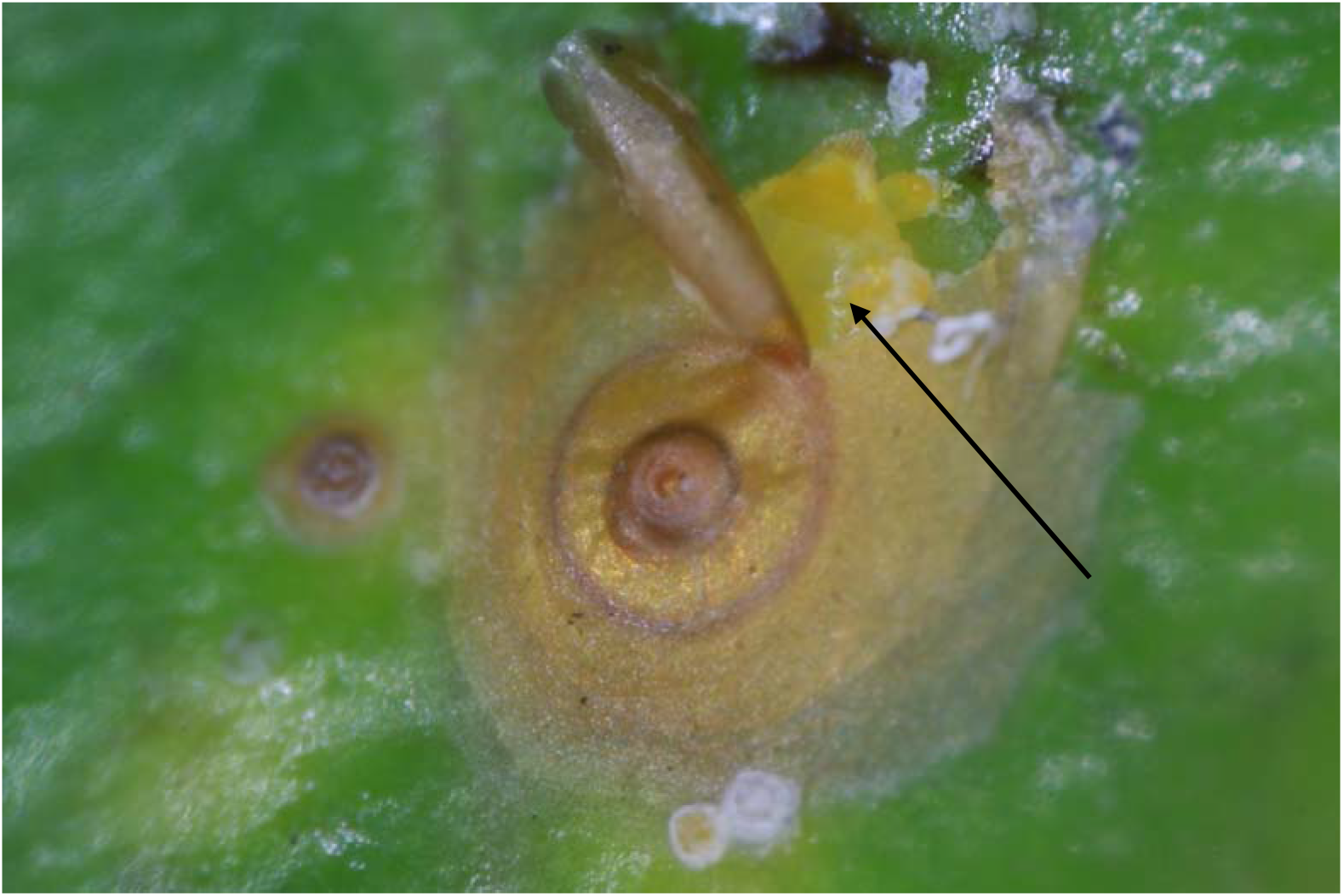
Body of female of *Chrysomphalus dictyospermi* (photo authors A.V. Shipulin, N.A. Gura)

Next, Table 3, Figure 3 shows an illustrative material from a microscopic examination of the pygidium of a brown shield female indicating the main diagnostic structures detected on the microscope slides.

**Table 3.** The main diagnostic micro-signs of the brown shield *Chrysomphalus dictyospermi* detected on the micropreparation slides.

**Fig. 3.**
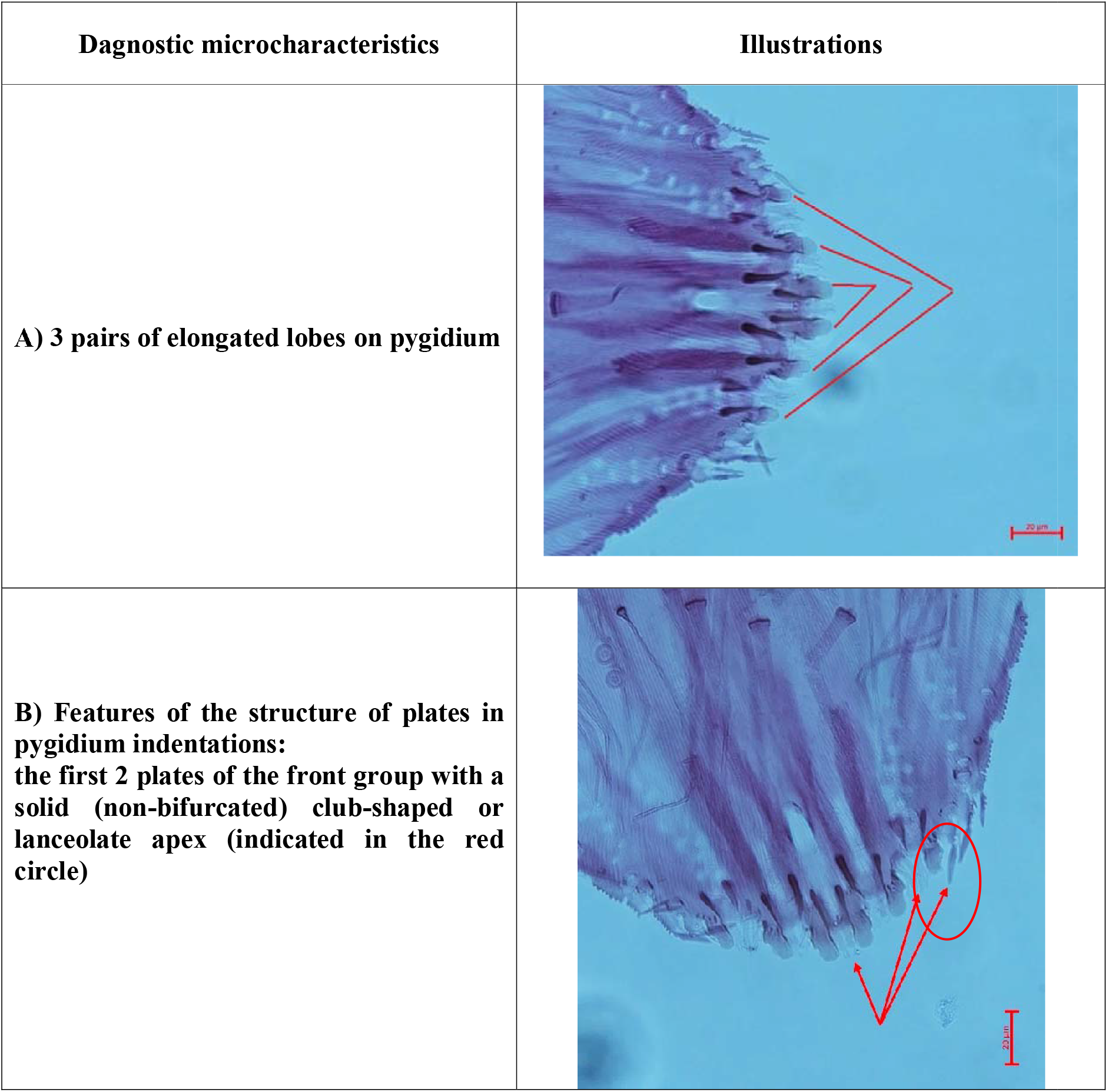
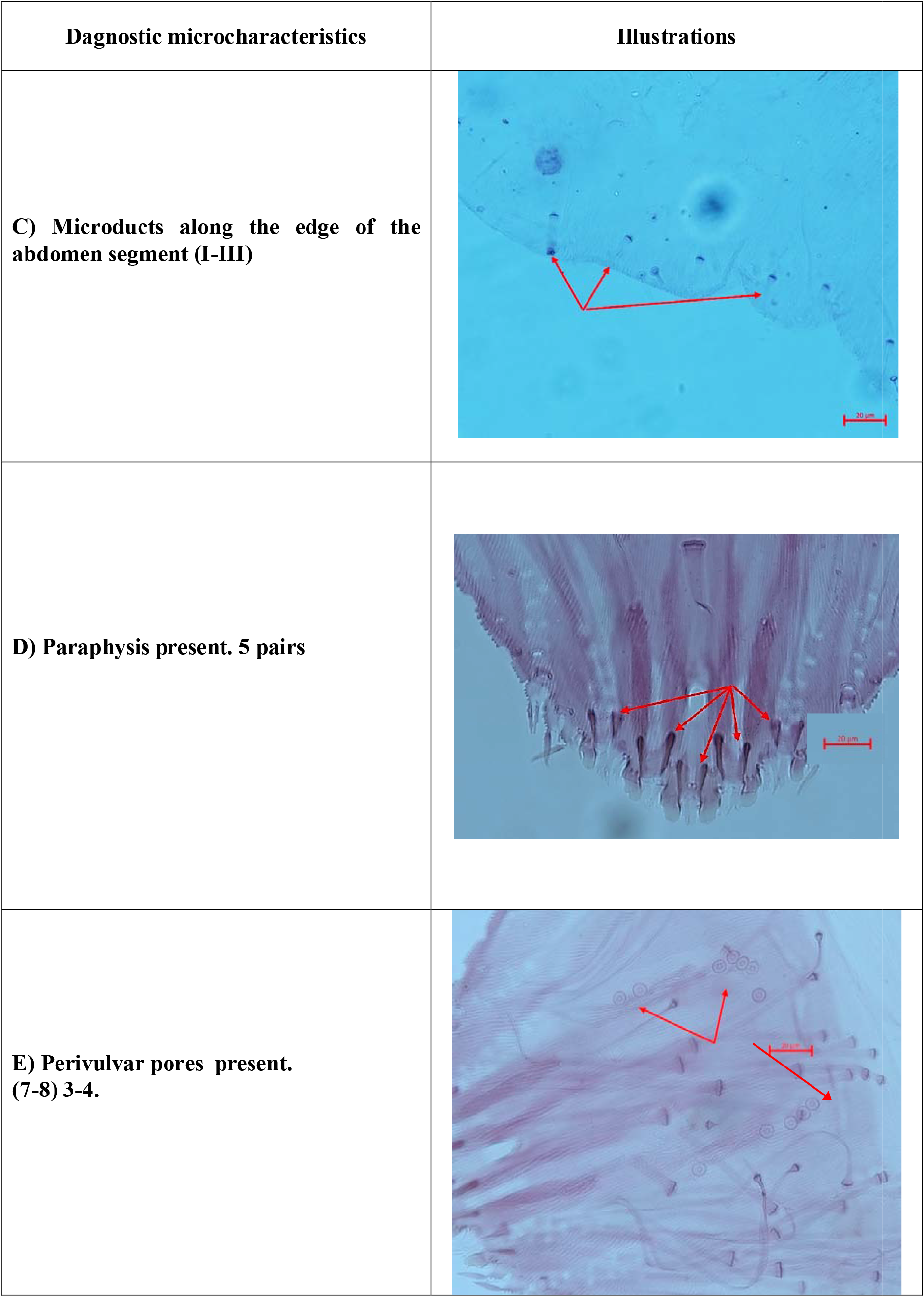
The main diagnostic micro-signs of the female *Chrysomphalus dictyospermi* (photo A-E - authors A.V. Shipulin, N.A. Gura)

Tables 4-5 show comparative analysis and illustrative material of diagnostic macro- and micro-signs of *Chrysomphalus dictyospermi* and closely related species.

**Table 4.**
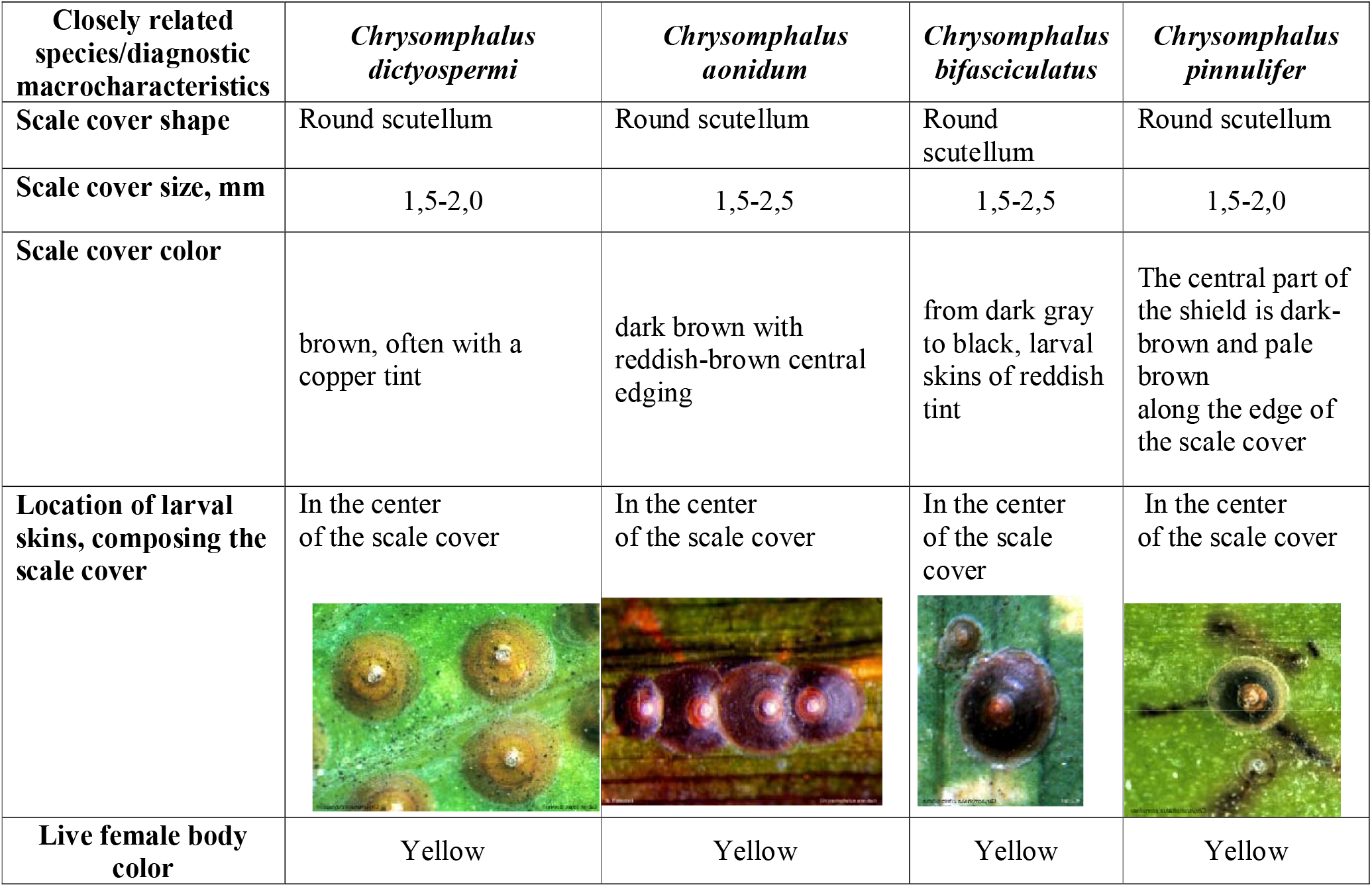
Comparative analysis of macrocharacteristics of closely related *Chrysomphalus* sp. (https://diaspididae.linnaeus.naturalis.nl/linnaeus_ng/app/views/introduction/topic.php?id=3377&epi=155)

| Closely related species/diagnostic macrocharacteristics | *Chrysomphalus dictyospermi* | *Chrysomphalus aonidum* | *Chrysomphalus bifasciculatus* | *Chrysomphalus pinnulifer* |
| --- | --- | --- | --- | --- |
| **Scale cover shape** | Round scutellum | Round scutellum | Round scutellum | Round scutellum |
| **Scale cover size, mm** | 1,5-2,0 | 1,5-2,5 | 1,5-2,5 | 1,5-2,0 |
| **Scale cover color** | brown, often with a copper tint | dark brown with reddish-brown central edging | from dark gray to black, larval skins of reddish tint | The central part of the shield is dark-brown and pale brown along the edge of the scale cover |
| **Location of larval skins, composing the scale cover** | In the center of the scale cover | In the center of the scale cover | In the center of the scale cover | In the center of the scale cover |
| **Live female body color** | Yellow | Yellow | Yellow | Yellow |

**Table 5.**
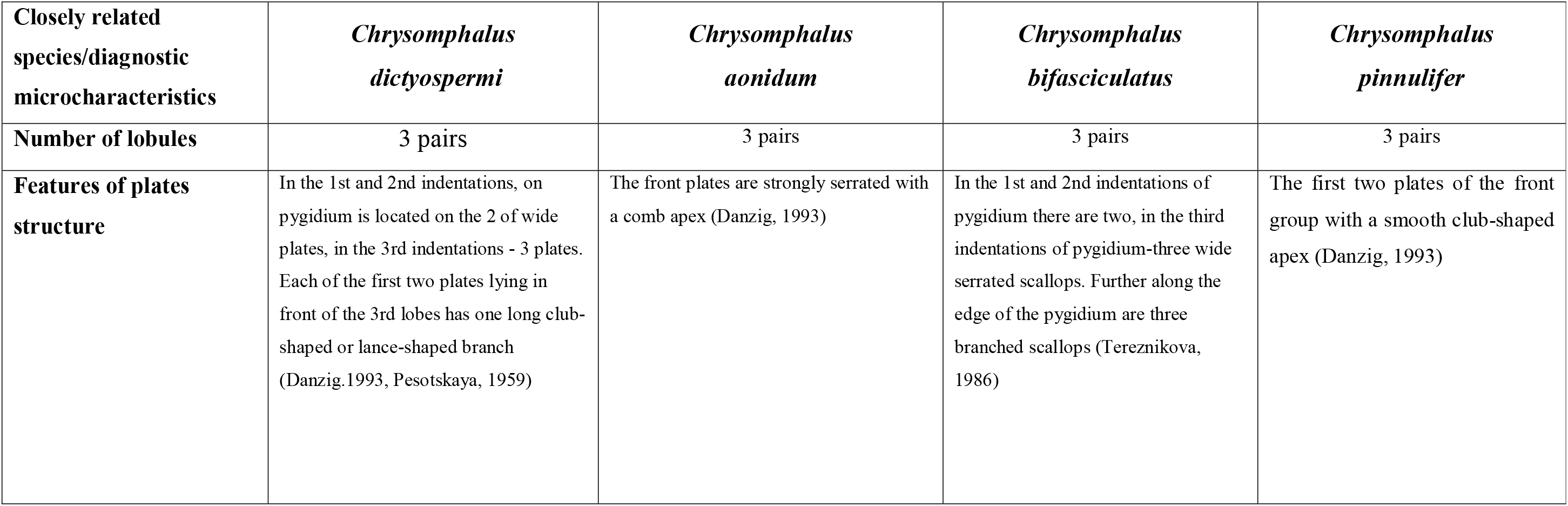
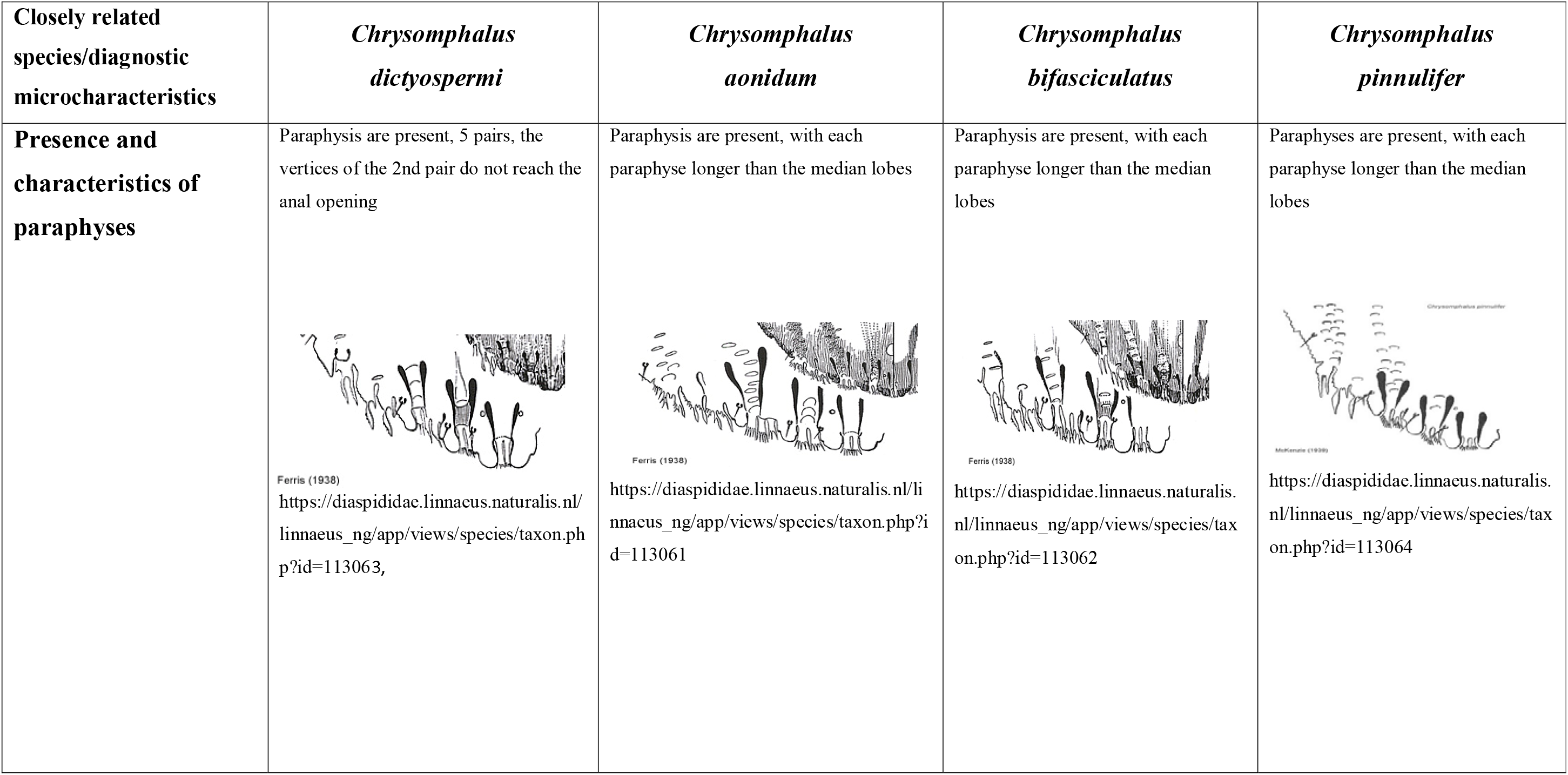
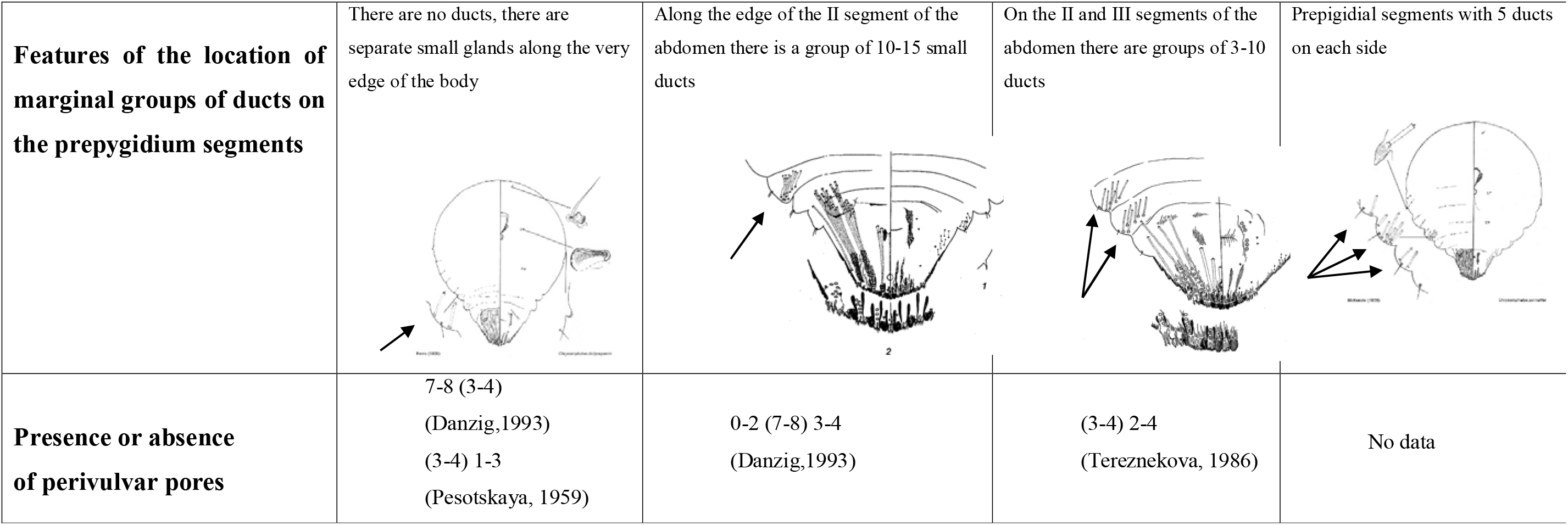
Comparative analysis of microcharacteristics of closely related *Chrysomphalus* sp. (https://diaspididae.linnaeus.naturalis.nl/linnaeus_ng/app/views/introduction/topic.php?id=3377&epi=155)

Among the main microcharacteristics of female pygidium of *Chrysomphalus* genus, the following have been analyzed: the number of lobes and their shape, the presence of paraphyses and their structure, the presence or absence of perivulvar pores. The results of comparative analysis of diagnostic microcharacteistics of four species of *Chrysomphalus* sp. scales are presented in Table 5.

## Conclusions

1. Based on the results of a study of the structure of the dictyospermum scale female pygidium, an illustrative material of the main diagnostic features identified on a microslide is presented.
2. Diagnostic macro- and microfeatures of four closely related species of *Chrysomphalus* genus are analyzed and given in the form of comparative tables, which allows us to reliably distinguish a quarantined object from an non-quarantine species of scales during laboratory entomological study, which is an important part of phytosanitary procedures. This paper can be used by quarantine and plant protection specialists involved in the diagnosis of detected organisms.

